# N-linked glycosylation increases horse radish peroxidase rigidity leading to enhanced activity and stability

**DOI:** 10.1101/2022.09.23.509167

**Authors:** Krithika Ramakrishnan, Rachel L. Johnson, Sam D. Winter, Harley L. Worthy, Chris Thomas, Diana Humer, Oliver Spadiut, Sarah H. Hindson, Stephen Wells, Andrew H. Barratt, Georgina E. Menzies, Christopher R. Pudney, D. Dafydd Jones

## Abstract

Glycosylation is the most prevalent protein post-translational modification, with a quarter of glycosylated proteins having enzymatic properties. Yet the full impact of glycosylation on the protein structure-function relationship, especially in enzymes, is still limited. Here we show glycosylation rigidifies the important commercial enzyme horseradish peroxidase (HRP), which in turn increases its activity and stability. Circular dichroism spectroscopy revealed that glycosylation increased holo-HRP’s thermal stability and promoted significant helical structure in the absence of haem (apo-HRP). Glycosylation also resulted in a 10-fold increase in enzymatic turnover towards *o*-phenylenediamine dihydrochloride when compared to its non-glycosylated form. Utilising a naturally occurring site-specific probe of active site flexibility (Trp117) in combination with red-edge excitation shift fluorescence spectroscopy, we found that glycosylation significantly rigidified the enzyme. *In silico* simulations confirmed that glycosylation largely decreased protein backbone flexibility, especially in regions close to the active site and the substrate access channel. Thus, our data shows that glycosylation does not just have a passive effect on HRP stability but can exert long range effects that mediate the ‘native’ enzyme’s activity and stability through changes in inherent dynamics.

## Introduction

Post-translational modification (PTM) of proteins is a common event in nature playing an essential role in imparting new structural and functional features on a protein^1^. Cofactors such as haem expands the chemistry available to proteins allowing them to perform functions ranging from O_2_ transport to electron transfer to catalysis^2^. Glycosylation is the most prevalent PTM, with N-linked glycosylation (attachment via an asparagine residue in N-X-S/T sequence motif) the most common attachment mechanism^3,4^. UniProt (www.uniprot.org/uniprotkb) reports that just over 87,000 proteins are glycosylated with *circa* 23,000 of those displaying catalytic activity. Given the number of glycosylated enzymes, it is important that we understand the impact of glycosylation on the enzyme structure-function relationship, especially if the enzymes are subsequently used for biotechnological applications and recombinant production is moved to different organisms.

While protein glycosylation is ubiquitous in eukaryotes, our knowledge of its impact on the protein structure-function relationship is still limited. Glycosylation plays several important biological roles^3^ including protein quality control and molecular recognition. Glycan addition can also stabilise proteins by increasing solubility, promoting correct folding, making them more resistant to proteolysis and preventing aggregation^3,5–7^. More recently, molecular dynamic simulations have suggested glycosylation can make a protein component less flexible^8^ despite the inherent flexibility of the glycans themselves^3,9^. The question arises is how glycosylation influences protein structure and thus in turn function, especially for enzymes where aspects such as dynamics can play a key role in defining catalysis^10,11^.

Using horseradish peroxidase (HRP) as a model enzyme system, we probed experimentally how glycosylation together with co-factor [haem] binding affects the dynamics and thus function and stability. HRP is an important commercial haem-dependent enzyme that has been extensively studied as a model for metal-dependent oxidation reactions^12–14^. HRP is naturally derived from the roots of the Horseradish plant, *Armoracia rusticana*, with different isoforms produced. The most common isoform is C1A^15^, which has in turn been the focus of recombinant expression^16,17^, structural studies^18, 19^, and protein engineering^14, 20, 21^. HRP is monomeric and binds haem non-covalently with His170 coordinating the haem iron ion (Figure 1). The naturally produced enzyme has eight N-linked glycosylation sites^22–25^. Recombinant versions produced in bacteria suggest that glycosylation is not absolutely essential to folding or activity of the enzyme.^16, 18^ Nevertheless, the absence of glycosylation has been reported to effect physicochemical properties of HRP such as folding efficiency, solubility and aggregation propensity.^26–28^

**Figure 1.**
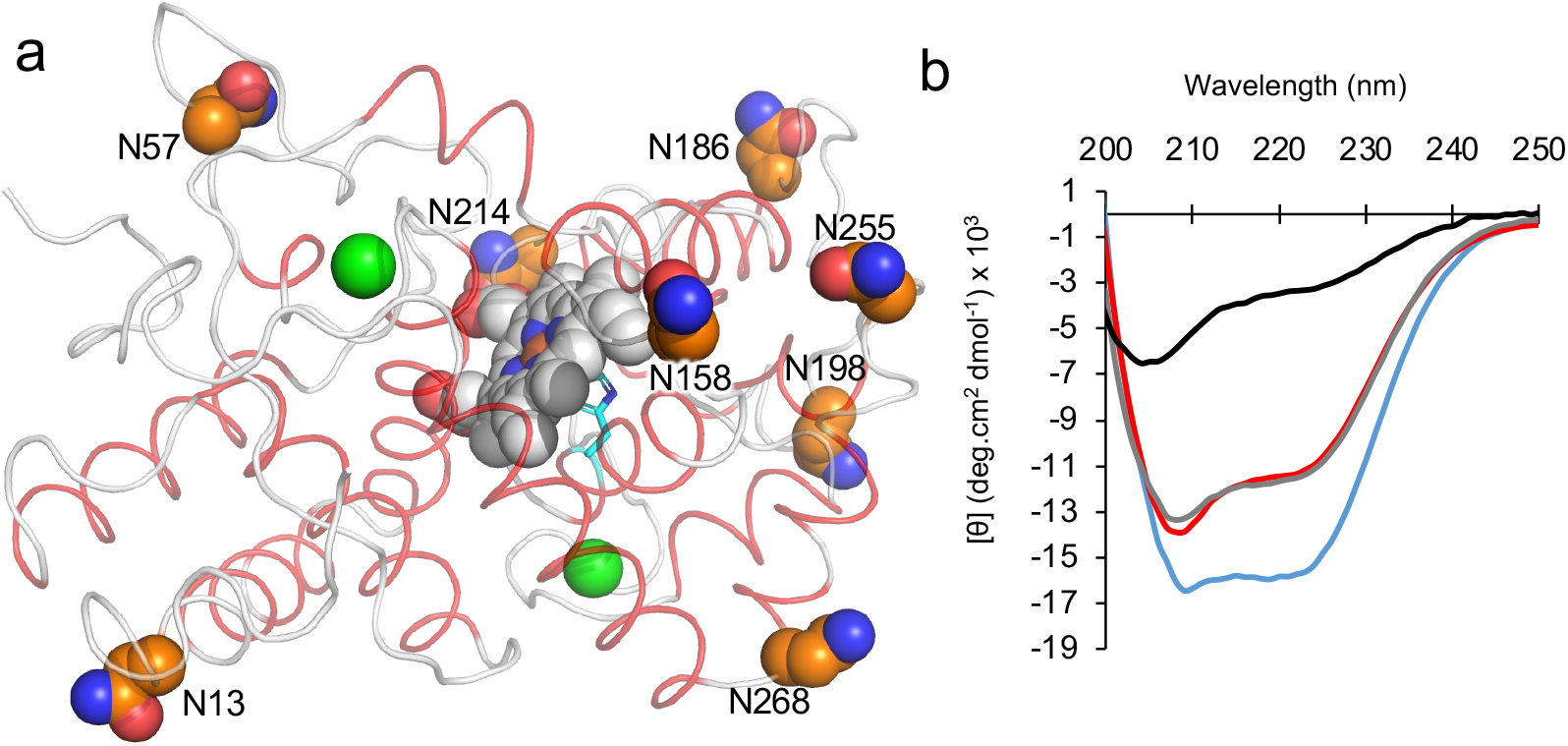
Structure of HRP. (a) Structure of HRP^19^(PDB 1hch). Helical regions are coloured red, haem is shown as grey spheres, with the iron co-ordinating H170 shown as cyan coloured sticks and calcium ion as green spheres. The N-linked glycosylated asparagine residues are shown as orange spheres. (b) Far-UV CD spectra of holo and apo version of the glycosylated plant-produced pHRP (blue and red, respectively) and the recombinantly produced holo- and apo-rHRP (grey and black, respectively) at 25° C.

Here, we find that the glycosylated holo-HRP is the most stable form of the enzyme and has a significantly higher catalytic turnover than the recombinant holo-HRP. Using a site-specific fluorescence probe close to the active site (117) naturally present in HRP, we experimentally find that the glycosylated holo-HRP has the most rigid active site but rigidification does not affect the temperature optimum for catalysis. Molecular dynamics and rigidity analysis reveals that glycosylation makes HRP less dynamic, especially around the active site. Thus, it appears that surface glycosylation not only globally stabilizes HRP but exerts long range rigidifying effects into the core of the enzyme that influences catalysis.

## Experimental methods

### Protein production

Lyophilized glycosylated apo and holo plant HRP was obtained from Ortho Clinical Diagnostics (Pencoed, UK). Haem occupancy was confirmed spectrophotometrically yielding a Reinheitszahl constant (RZ A_404_/A_280_) 2.98 for holo protein (Figure S2). Apo-pHRP was as generated by Biozyme for Ortho Clinical Diagnostics and manufactured using a butanone extraction approach^29^ before buffer exchange into water and lyophilisation. The composition of the pHRP protein was determined by ESI-ToF Mass Spectrometry performed by the Analytical Services facility within the School of Chemistry, Cardiff University. Recombinant holo HRP was produced as described previously^17^. Haem occupancy was confirmed spectrophotometrically yielding a Reinheitszahl constant (RZ A_404_/A_280_) 2.91 (Figure S2). Haem was removed using a butanone extraction approach as described previously^29^.

### Circular Dichroism

Circular Dichroism measurements were performed on a Chirascan TM using 5-10 μM protein in either 20 mM Bis-Tris (pH 7.0) for rHRP or 50 mM Tris (pH 8.0) for pHRP. Spectra were recorded from 200-260 nm at 1 nm intervals. Samples were subsequently heated at a ramp rate of 1°C per minute using a Quantum Northwest Peltier with absorbance at 222 nm at every degree change up to 90°C. The subsequent melting curves were fit a Boltzmann sigmoidal two-state transition model (Figure 2a),

**Figure 2.**
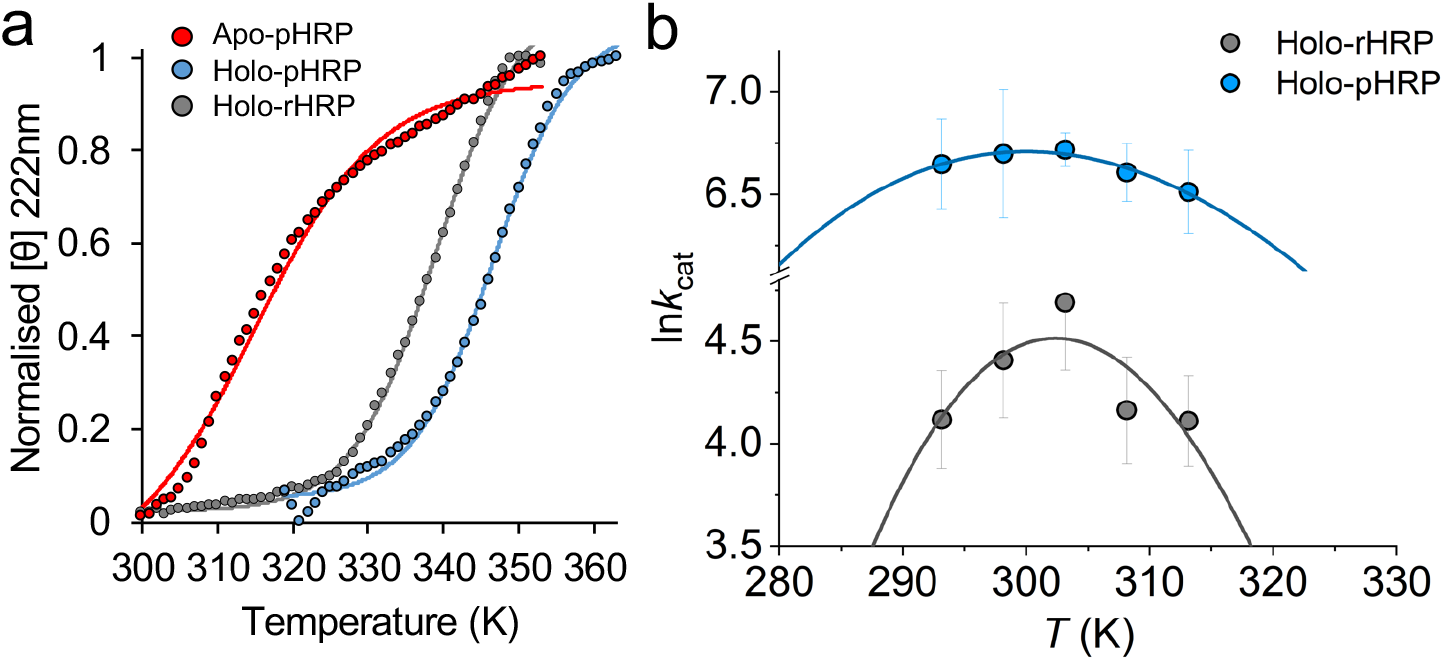
Temperature dependence of HRP structure and function. (a) Thermal melts measured by CD at 222 nm. The normalised molar ellipticity ([Θ]) was calculated by setting the value at 300 K to 0 and the value at 353 K (apo-pHRP and holo-rHRP) or 363 K (holo-pHRP) to 1. The full thermal melt data is presented in Supporting Figure S3, together with the data for apo-rHRP that shows no clear transition. The dashed line is the Boltzmann sigmoidal fit to a two-state transition. (b) Temperature dependence of HRP turnover. Solid lines are the fits to Eq 1. Apo-pHRP is shown as red lines, holo-pHRP as blue lines and holo-rHRP as grey lines.

### Steady State Kinetics

Steady state kinetics of HRP was carried out using the substrate *o*-phenylenediamine dihydrochloride (OPD) (Sigma Aldrich) that is converted to the coloured product 2,3-diaminophenazine (DAP)^30,31^. For rHRP, the reaction conditions were 20 mM Bis-Tris (pH 7.0), 3% v/v hydrogen peroxide and 10 nM enzyme. For pHRP the reaction conditions were 50 mM Tris (pH 8.0), 3% v/v hydrogen peroxide and 0.5 nM enzyme. Protein concentrations were calculated using 403 nm absorbance (ε = 100 mM^-1^cm^-1^) so only the concentration of holo-protein is considered. Freshly prepared OPD substrate ranging in concentration from 0.1 mM to 4.0 mM was added to the reaction mix. Absorbance was recorded at 450 nm over 60 seconds on a Cary UV Spectrophotometer, in triplets for each substrate concentration. The substrate turnover per minute was calculated from the absorbance change over 1 minute using the extinction coefficient of DAP product at 450 nm (10,600 M^-1^cm^-1^). The kinetic data was fitted to a Michaelis-Menten equation using GraphPad Prism software to determine V_max_, *K*_M_, and *k*_cat_. Reaction rates of pHRP in 20 mM Bis-Tris (pH 7.0) where comparable to those in 50 mM Tris (pH 8.0). However, rHRP rates were lower in 50 mM Tris (pH 8.0) thus we used the more optimal 20 mM Bis-Tris (pH 7.0) buffering conditions to measure rHRP kinetics.

Temperature dependent kinetics data were also obtained on a Cary UV Spectrophotometer where all reaction components were incubated at temperatures ranging from 293K (20°C) to 313K (40°C) for 1 hour before mixing the components and recording the absorbance at 450 nm. The substrate (OPD) concentration used was equal to 10x of *K*_M_ value and absorbance change was recorded at 450 nm for 2 minutes, in triplicates for each temperature. Substrate turnover per minute was calculated as previously described using the extinction coefficient of DAP product.

### Red Edge Emission Shift

Red edge emission shift (REES) of HRP was determined by measurement of fluorescence emission using 5 μM protein samples which were prepared under the buffer conditions used for the steady-state kinetics. Proteins were excited at 1 nm intervals over the range 292 nm to 310 nm and emission was recorded between 315 nm to 500 nm at a scan rate of 30 nm/min. For all readings, excitation and emission slit width was set to 5 nm with a detector voltage set to high. Up to 12 independent readings at each excitation wavelength were taken and the average calculated.

The quantification of the REES data relies on accurate extraction of information on how the structure of the emission spectra vary and so the additional band would convolve the measurement.

As in our previous work^32–35^, we therefore numerically modelled each of the spectra using a sum of two skewed Gaussians (Eq 1) as described recently for a *de novo* haem peroxidase^33^

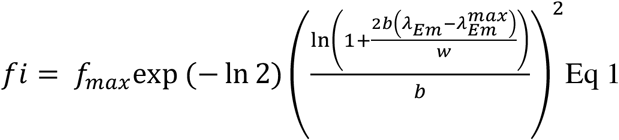

where *fi* is the measured fluorescence intensity, *f*_max_ is the maximum emission intensity at wavelength 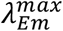, with a full width at half maximal of w and the ‘skewness’ was controlled by *b*. Fluorescence spectra were accurately modelled and deconvolved by fitting to such functions as demonstrated by elsewhere.^32–35^ By fitting to a sum of two skewed Gaussians we were able to accurately model the spectral component attributable to tryptophan emission alone. From these models the centre of spectral mass (CSM) was extracted for each spectral component, which allowed quantification of changes in the structure of the fluorescence spectra,

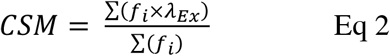

The resulting plot of CSM versus the change in excitation wavelength, Δ*λ_Ex_*, is shown in Figure 2a for each of the HRP versions.

### Molecular dynamics

The PDB model for HRP (1h58) was uploaded to the solution builder page on the CHARMM-GUI server (^36^, https://www.charmm-gui.org/?doc=input/solution) transferring the coordinates for the protein, heme residue and calcium ions. At the protein modification stage, default disulphide bonds were retained, and a heme coordination site was added at residue H170. To generate the glycosylated form of the protein, (Xyl)Man3(Fuc)GlcNac2 - N-1 (see figure S5) was added to asparagine residues: N13, N57, N158, N186, N198, N214, N255, and N268 at the PDB manipulation stage of the CHARMM-GUI solution builder. The simulations were set up in a cubic box with a 10 nm edge distance. Protein charge was neutralised by the addition of 150 mM CaCl_2_ ions and solvated. PME FFT grid was generated automatically by the server. Parameters and input files for use in GROMACS using the CHARMM36m forcefield, were then generated with hydrogen mass repartitioning and WYF cation-pi interactions^37,38^. Input files were downloaded from the server to run molecular dynamics (MD). MD was run through three stages, minimisation, equilibration and production. Minimisation used the steepest decent method with a tolerance of 1000 KJ^-1^ nm^-1^ and a cut off-off 5000 steps. The equilibrium stage was run for 125000 steps with a 0.001 fs time per step. Finally, the production run was carried out for 100 ns total time with a 0.004 fs time per step. A temperature of 303.15°K, a pressure of 1 atm and periodic boundary conditions were applied to all simulations. For all simulations, Nose-Hoover temperature coupling thermostat was applied and Particle-mesh Ewald (PME) was applied to long range electrostatics. Analysis was carried out on both production runs using inbuilt GROMACS modules.

### Pairwise network analysis

The Ca-Ca distance changes were determined as follows. Using the GROMACS mdmat command, a distance matrix was generated for both non-glycosylated and glycosylated MD simulations which consisted of the smallest distance between each residue over the course of each simulation. This distance matrix was converted from xpm to csv format using a python script. Both distance matrices were then parsed using a custom jupyter notebook and merged to calculate the change in Cα-Cα distance. A function was written to only consider the change in Cα-Cα distances between residues that were within 5.5Å. Another function was written to only consider Cα-Cα distances between residues which were greater than i+4 apart. The resulting adjacency list was analysed and visualised as a network using Cytoscape 3.9.1^39^. The interactions networks was determined as follows. Using PyMol, GROMACS trajectories were converted to PDB ensembles. Resulting PDB ensembles were uploaded to the RING 3.0^40^ server to identify non-covalent interactions and their frequency over the duration of the simulation. “Strict” distance thresholds were used to classify inter-residue interactions. The resulting machine-readable JSON files were converted to csv using Cytoscape. Both csv files were then merged using a custom jupyter notebook where a function was written to only extract interaction frequencies that underwent a change +-10%. The resulting csv file was imported to Cytoscape for analysis and network visualisation. All scripts and notebooks used in this analysis are available at https://github.com/DrewBarratt/MD_Network_analysis

### Rigidity analysis

Pebble-game rigidity analysis^41, 42^ is an integer algorithm which can divide a protein structure into rigid clusters and flexible regions by matching degrees of freedom against bonding constraints on a directed graph constructed from the covalent and noncovalent interactions of the protein. Glycosylation was modelled by the addition of artificial constraints to the covalent interaction network, reducing the conformational flexibility of the backbone at glycosylation sites. The results of this division, or Rigid Cluster Decomposition (RCD), depend on which noncovalent interactions are included, and are therefore a function of a hydrogen bond energy cut-off which excludes weaker polar interactions such as hydrogen bonds. A “rigidity dilution” is carried out by progressively altering this cut-off from small negative values to larger negative values, gradually excluding polar interactions from weaker to stronger. The most rigid portions of the structure can be identified as those which retain rigidity longest during this dilution. The analysis was carried out on an all-atom model of the protein and rigid clusters can extend across main-chain and sidechain structural groups. For this study, a residue is considered part of the largest rigid cluster (RC1) if its Ca is part of that cluster. The rigidity analysis software used here (“FLEXOME”, written by SAW) is available on request from the University of Bath (https://doi.org/10.15125/BATH-00940).

## Results

### The effect of glycosylation and haem on HRP stability

Two sources of HRP were used here: directly from the plant (referred from herein as plant or pHRP) and a recombinant source produced in *E. coli* (referred from herein as recombinant or rHRP). The rHRP is based on the C1A isoform and its production has been described previously^17^. The pHRP is commonly sourced and utilised by companies, including diagnostic companies; our source is clinical diagnostic grade pHRP used in tests manufactured by OrthoClinical Diagnostics. The pHRP is produced in the plant as the glycosylated holo-form with haem bound. The apo form of pHRP is subsequently generated by means of a commonly used denaturation-organic extraction process that removes non-covalently bound haem^29^. In immunoassays, apo-pHRP is used as a correction agent for false positives. As we want to directly compare our pHRP with a defined recombinant HRP source, we used mass spectrometry to confirm the composition of both pHRP forms (Supporting Figure S1, Supporting analysis). Apo- and holo-pHRP contain the same species that match the commonly observed C1A (with and without the C-terminal serine) glycosylated forms^23,43^. The absorbance spectra of pHRP and rHRP are very similar with a λ_max_ at 402-404 nm (Figure S2). Thus, the main difference between pHRP and rHRP is the presence of N-linked glycans so allowing us to investigate the effects of glycosylation on the enzyme and how glycosylation affects the structure and stability of the apo-form.

We first looked at the impact of each PTM event on general structure and stability of HRP using far-UV circular dichroism (CD) spectroscopy. All HRP forms apart from apo-rHRP show far-UV CD spectra at 25°C that are characteristic of predominately α-helical structure reflective of the helical nature of HRP (Figure 1b), exhibiting two signature minima at ~210 and ~220 nm (Figure 1b). Glycosylated holo-pHRP has deepest 210/222 nm troughs suggesting it has the highest helical content. The apo-rHRP spectrum shows the protein is largely unfolded confirming the importance of haem to overall structural integrity. However, glycosylation can compensate for the loss of haem with the apo-pHRP having a similar spectral profile to holo-rHRP. These data therefore suggest that, at least in terms of the protein’s secondary structure formation, HRP’s global structure is influenced by both haem binding and glycosylation, with the latter promoting helical structure in the apo-form of HRP as well as the functional holo-protein.

Thermal unfolding was then measured to probe the effect of each PTM on stability. Apart from apo-rHRP, the CD thermal melts (Figure 2a) conform to apparently simple two-state unfolding transitions. Our data suggest both haem binding and glycosylation stabilize HRP structure, with haem having the most significant effect. Holo-pHRP is the most thermally stable with a Tm of 348 K, 10 K higher than holo-rHRP (Tm 338 K). Apo-pHRP melts at a lower temperature than both holo protein forms (T_m_ 314 K), highlighting the integral nature of co-factor binding to structural stability. Apo-rHRP has a constant 222 nm signal with no clear transition confirming that it is largely unfolded prior to temperature ramping (Supporting Figure S3). Thus, there is a clear stabilizing effect of haem binding, translating to a > 30°C stabilisation (in terms of T_m_). Glycosylation is also having a significant effect by increasing the stability of the holo protein and promoting the folding of the apo-form, or at the very least promoting and/or stabilizing helical elements. The effect of glycosylation is also consistent between the holo and apo enzyme, giving an increase in T_m_ on glycosylation of at least 10°C. These observations point to a synergistic benefit of the combination of glycosylation and haem binding on HRP thermal stability.

### The effect of glycosylation on enzyme turnover

We next explored the effect of glycosylation on the steady-state kinetics of HRP and the effect on the temperature dependence of HRP turnover by monitoring the conversion of the commonly used substrate *o*-phenylenediamine (OPD) to the coloured 2,3-diaminophenazine (DAP)^30, 31^. Table 1 shows the Michaelis-Menten kinetics for holo-rHRP and glycosylated holo-pHRP at 298 K. The most significant effect of glycosylation is the large increase in turnover, with *k*_cat_ being an order of magnitude higher for holo-pHRP *versus* holo-rHRP. *K*_M_ remains relatively constant suggesting that it is catalysis rather than substrate binding being affected. Thus, glycosylation increases the rate of by which HRP turns over OPD (Table 1). The temperature dependence of *k*_cat_ for both holo-pHRP and holo-rHRP show significant curvature with respect to temperature (Figure 2b). Curvature in temperature dependence plots can arise from several sources, most commonly sub-saturation of substrate binding, protein unfolding and change in rate-limiting step. In the absence of these factors the curvature in temperature dependencies can reflect useful information on the thermodynamics of the chemical step and can be fit

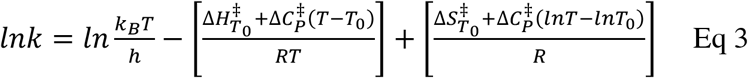

where *T*_0_ is an arbitrary reference temperature. 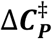 is the difference in heat capacity between the ground and transition states. 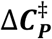 determines the change in ΔH^‡^ and ΔS^‡^ with temperature and thereby defines the non-linearity of the temperature dependence of the Gibbs free energy difference between the ground state and the transition state (ΔG^‡^).

**Table 1.**
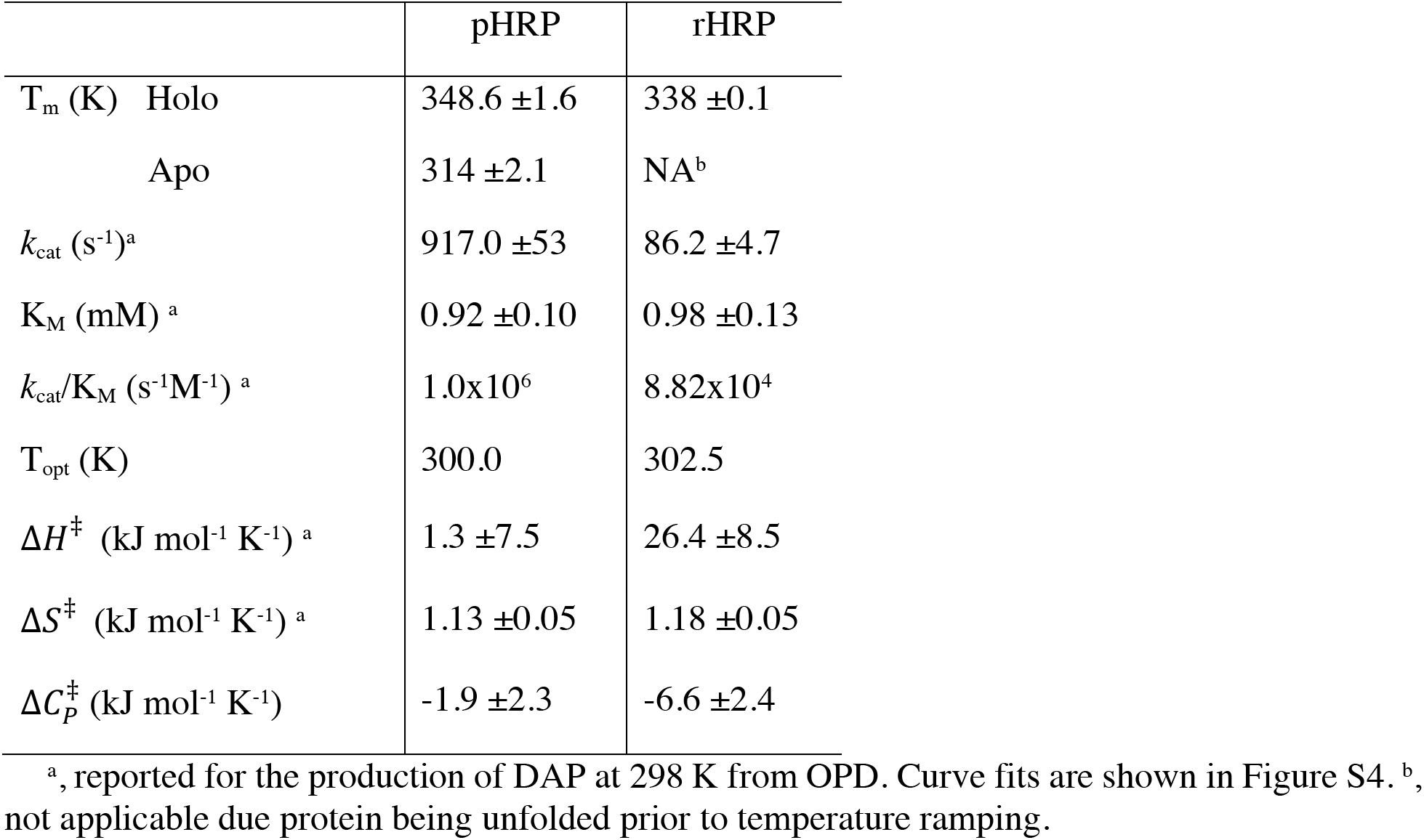
Thermodynamic and kinetic parameters of holo HRP.

|  | pHRP | rHRP |
| --- | --- | --- |
| $T_m$ (K) Holo | 348.6 ± 1.6 | 338 ± 0.1 |
| Apo | 314 ± 2.1 | NA<sup>b</sup> |
| $k_{cat}$ (s<sup>-1</sup>)<sup>a</sup> | 917.0 ± 53 | 86.2 ± 4.7 |
| $K_M$ (mM) <sup>a</sup> | 0.92 ± 0.10 | 0.98 ± 0.13 |
| $k_{cat}/K_M$ (s<sup>-1</sup>M<sup>-1</sup>) <sup>a</sup> | 1.0×10<sup>6</sup> | 8.82×10<sup>4</sup> |
| $T_{opt}$ (K) | 300.0 | 302.5 |
| $\Delta H^\ddagger$ (kJ mol<sup>-1</sup> K<sup>-1</sup>) <sup>a</sup> | 1.3 ± 7.5 | 26.4 ± 8.5 |
| $\Delta S^\ddagger$ (kJ mol<sup>-1</sup> K<sup>-1</sup>) <sup>a</sup> | 1.13 ± 0.05 | 1.18 ± 0.05 |
| $\Delta C_P^\ddagger$ (kJ mol<sup>-1</sup> K<sup>-1</sup>) | -1.9 ± 2.3 | -6.6 ± 2.4 |

<sup>a</sup>, reported for the production of DAP at 298 K from OPD. Curve fits are shown in Figure S4. <sup>b</sup>, not applicable due protein being unfolded prior to temperature ramping.

We note our temperature-dependent assays are performed with an excess of substrate (10x *K*_M_) and so the curvature is not due to sub-saturation of the *K*_m_ at elevated temperatures. Moreover, our steady-state progress curves are entirely linear over the time course of the measurement, suggesting that there is no appreciable unfolding of HRP during the assay as the *T*_m_ values from CD thermal melts are ~30 °C higher than the highest temperature used in our kinetic studies. Therefore, fitting using the MMRT model appears appropriate. We cannot however explicitly rule out a change in rate limiting step with respect to temperature and so we focus our analysis on the apparent temperature optimum, rather than the microscopic interpretation of the magnitude of 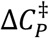. The parameters resulting from fits to the temperature dependence of *k*_cat_ are given in Table 1; the only significant difference in the two HRP forms arises from the increased rate constant for pHRP over rHRP (Figure 2a). Indeed, despite the measurable (Δ*T*_M_ ~ 10°C; Table 1) difference in thermal stability and turnover number, the temperature optimum for enzyme turnover is essentially identical. That is, the temperature optimum for catalysis is not linked to the thermal stability of the enzyme.

### Using fluorescence to probe the effect of glycosylation on HRP rigidity

Given that both enzyme kinetics and thermal stability improved on glycosylation of HRP (Table 1), we experimentally investigated the local effect of glycosylation on active site rigidity/flexibility. As shown in Figure 1a, all 8 surface-exposed N-linked glycosylation sites are distant from the largely buried active site; the closest N-linked site to the haem centre is N214 at ~20Å. We have recently developed the use of an optical phenomenon that allows changes in the flexibility of a protein to be assessed via tryptophan fluorescence measurements: the red edge excitation shift (REES)^11,32,34,44^. REES is an optical phenomenon which tracks changes in the distribution of protein conformational states, via shifts in solvent-fluorophore [Trp] interaction energies^45^. We find that the REES phenomenon, when quantified using our approach (described below), provides an extraordinarily sensitive metric of changes in flexibility. HRP affords an ideal, natural, site-specific probe of the active site volume/rigidity as it has a single tryptophan residue (Trp117), 9 Å from the haem (Figure 3a). We have recently applied REES to engineered porphyrin binding proteins^33,46^ and were able to discriminate between differently flexible forms of a *de novo* designed artificial haem peroxidase that mapped precisely with NMR observations^33^.

**Figure 3.**
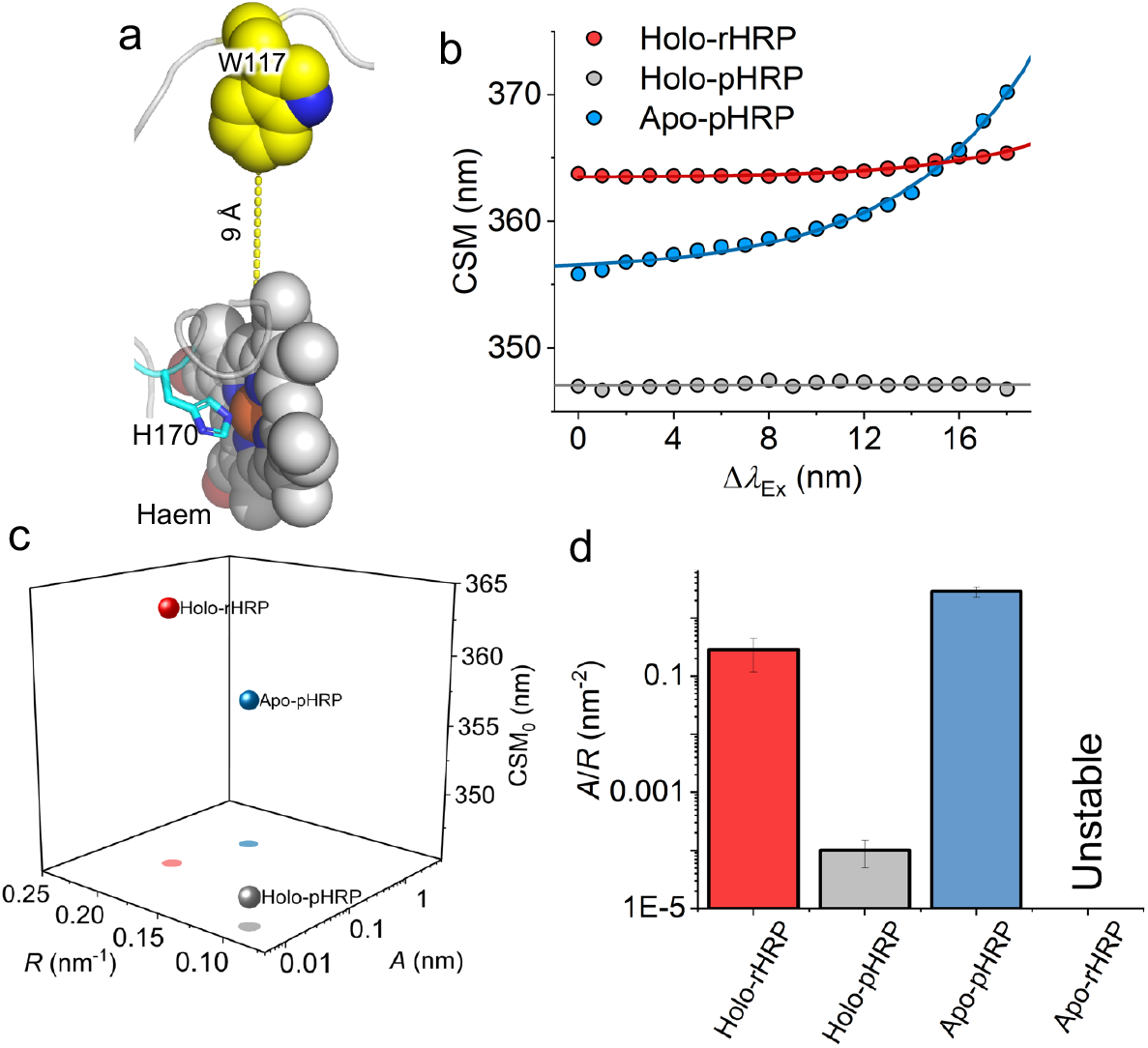
An active site-specific REES probe captures shifts in flexibility. (a) Location of Trp117 relative to haem. (b) Raw REES data, solid line shows the fit to Eq 4 as described in the main text. (c) Plots of parameters resulting from the fits shown in panel b. (d) Ratio of parameters used to reflect shifts in molecular flexibility.

Figure 3 shows the REES data analysis for each of the HRP versions studied, except for apo-rHRP; consistent with the findings from our CD data (Figure 1b), apo-rHRP showed a propensity to aggregate which significantly convoluted the fluorescence spectra. The REES data shows an upward curvature in the magnitude of centre of spectral mass (CSM; Figure 3b) with respect to change in emission wavelength (Δ*λ_Ex_*); for a single tryptophan containing protein, such curvature is indicative of a measurable REES effect and that the tryptophan is able to sample a range of different environments.

It is common to use 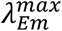 of tryptophan fluorescence spectra to report on changes in the solvent exposure of tryptophan residues, with an increase in solvent exposure leading to an increase in 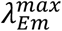. The parameter CSM_0_ therefore reports similarly but is more accurate since it accounts for excitation energy dependent changes in fluorescence spectra. To quantify the REES phenomenon so as to compare changes in flexibility we fit the REES data to a simple exponential function

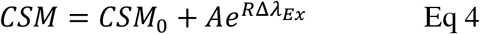

where the amplitude and curvature of the exponential is described by *A* and *R* respectively and CSM_0_ is the CSM value independent of λ_Ex_. The plot of the resulting values from these fits are then shown in Figure 2b. We have previously found that comparing the ratio of the *A* and *R* parameters is a simple way to infer changes in flexibility, with a large *A*/*R* value reflecting a more flexible protein/environment and smaller value reflecting a more rigid protein/environment.^11,32,44,47^

The data in Figure 3 shows that glycosylation (holo-pHRP *versus* holo-rHRP) results in a significant reduction in *A*/*R* and in CSM0, respectively. That is, glycosylation of the protein surface alters the rigidity of the partially buried active site. On removal of the haem (Apo-pHRP), we find that the *A*/*R* increases dramatically, as well as showing an increase in CSM0 compared to holo-pHRP; removal of the haem increases active site flexibility and solvent exposure. The influence of haem is not unexpected given it is buried within the protein and forms an integral structural component. The haem effect thus seems logical. The active site rigidification effect of glycosylation is unexpected as, compared to haem, the N-linked sites are relatively distant from the Trp117 probes site and surface exposed (Figure 1a). Thus, the REES analysis suggests that the active site becomes more rigid on both glycosylation and haem binding, with holo-pHRP being the most rigid.

### Molecular simulations of HRP dynamics

To further explore how glycosylation impacts on the dynamics of HRP, we undertook *in silico* simulations. There is currently no structure of the glycosylated HRP so glycan units were added *in silico* to each N-linked site. Based on our mass spectrometry data (Figure S1), the major holo-HRP species contains 8 N-linked glycan units (GlcNAc-2, Man-3, Fuc-1, Xyl-1)^23, 43^. The glycan units, shown in Supporting Figure S5, were added to the HRP structure using the PDB manipulation modules of the CHARMM-GUI solution builder. Molecular dynamics simulations for the glycosylated and non-glycosylated HRP (PDB entry 1HCH^19^ modified for use with GROMACS) were run for 100 ns (Figure S6).

Glycosylation on the whole reduces Ca flexibility, which is indicative of increased backbone rigidity, with only a few residues where flexibility increased (Figure 4a-b). This is in line with recent simulations of HRP with non-native glycans attached^48^. Of 306 residues, 126 residues have a significantly (>10% in the change in root-mean square fluctuation’s (ΔRMSF) maximal value) lower Ca RMSF on glycosylation while only 20 residues are significantly more flexible on glycosylation. Of the 8 N-linked asparagine residues, 5 show significant reduction in RMSF (N13, N57, N186, N198 and N255, N268) when glycosylated, with none exhibiting an increase; N255 underwent the largest relative drop on glycosylation (1.53 Å). The haem group was relatively stable, with only 0.07 Å RMSF difference between the two HRP forms. The imidazole side chain of the haem coordinating residue H170 is less flexible (ΔRMSF 0.26 Å) in the glycosylated form. The central probe for REES, Trp117, does not change significantly on glycosylation (Figure 4a) suggesting it is reporting on HRP dynamics as a whole rather than changes to the residue itself.

**Figure 4.**
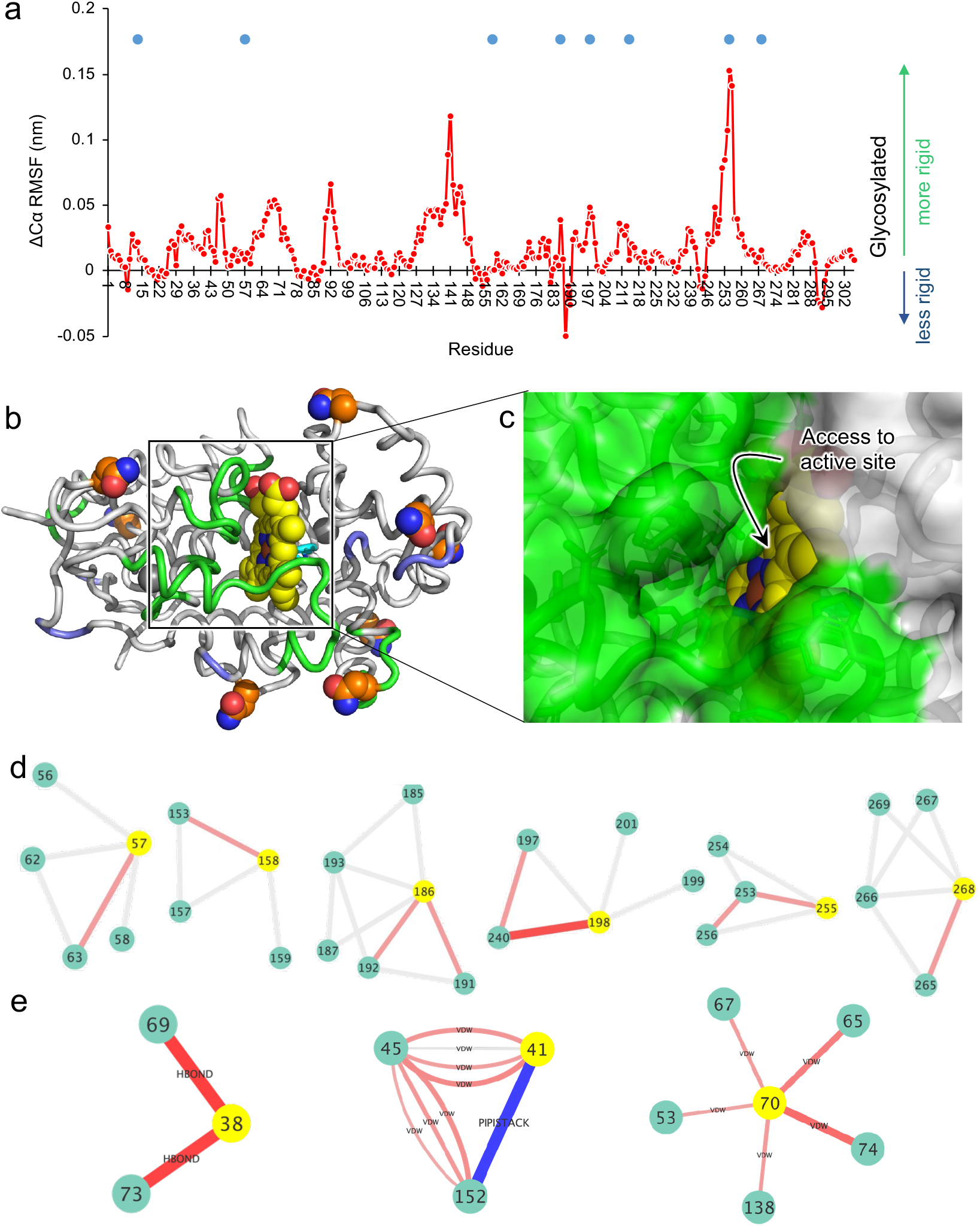
Molecular dynamic simulations of holo-HRP. (a) Change in root mean squared fluctuation (RMSF) for residue’s Ca over 100 ns. The blue circles represent the N-linked glycosylation sites. The actual Ca RMSF values for each form are shown in Figure S6b. The ΔCα was calculated by subtracting the RMSF value for the glycosylated holo-HRP from the unglycosylated form. (b) Mapping the RMSF changes onto the structure of HRP. Green regions become less dynamic on glycosylation and blue regions become more dynamic. N-linked sites are shown as orange spheres and haem as yellow spheres. (c) Surface representation of HRP showing the active site tunnel and access to the distal haem plane. Colours are as in (b). (d) Pairwise network for Ca atoms related to the N-linked sites (shown as yellow spheres) over the course of the MD simulation. The full Ca pairwise network is shown in Supporting Figure S7. Grey links represent no change in the Ca pairwise distances with red lines indicating closer distances with shade of red and thickness of the line related to the change in distance (ranging from light red thinner lines representing 0.38Å to thicker darker red lines representing 0.76Å). (e) Pairwise network of interactions for key catalytic residues (shown as yellow spheres) over the course of the simulation. Red lines indicate increased number of interactions in the glycosylated HRP and blue lines increased number of interactions in the non-glycosylated form. The thickness of the lines corresponds to the frequency of an interaction over the course of the MD, with thicker lines representing more persistent interactions. An arbitrary 10% cut off was applied, with interactions differences below this value ignored. The change in interaction type is shown on the diagram with VDW, HBOND and PIPISTACK equivalent to van der Waals, H-bonds and pi-pi stacking, respectively.

Mapping the significant changes on Ca RMSF onto the structure of HRP reveals that many of the regions that exhibited reduced flexibility lie close to the active site entrance and directly interacting with the distal, catalytic plane of haem (Figure 4b-c). Residues 68-72 together with residues 137-143 contribute towards forming the tunnel towards the central iron ion of haem and the substrate binding site. The latter region includes the conserved P139-A140-P141 motif commonly found in plant peroxidases^13^. Residues 248-257 undergo the largest drop in Ca RMSF on glycosylation and are located proximal to the active site tunnel residues. The clustered model from the glycosylated MD run suggests that glycan unit attached to N255 is potentially interacting with the N158 glycan unit. F41 together with H42 and its H-bond coupled partner, N70, are critical to catalysis^49^, with all three showing reduced Ca flexibility on glycosylation despite being buried and far from any of the N-linked glycosylation sites. H42 in particular is critical to catalysis as it plays an essential role in information of compound I through accepting a proton from H2O2. The region on either side of M284, which makes direct contact with haem, also becomes less flexible on glycosylation.

We then undertook a pair-wise network analysis to assess changes in the Ca distances over the course of the simulation, with a cut-off distance of 5.5 Å. While little change was observed for most pairs, significantly more Ca pairs were closer together in the glycosylated (64/614) form compared to the non-glycosylated form (26/614) (see Supporting Figure 8 for full pair-wise network). Moreover, a larger number of long range Ca pairwise distances (>i,i+4) were closer in the glycosylated HRP (42%) compared to the non-glycosylated enzyme (19%). Amongst the residues whose Ca become closer to their neighbours include key catalytic residues H170 (proximal haem iron coordination) and its associated residue N247 along with N70 (Supplementary Figure S7). With respect to the N-linked site, 6 of the 8 Asn Ca are closer to their neighbours on glycosylation (Figure 4d); the remaining two N-linked Asn sites (13 and 214) did not change significantly on glycosylation. This suggests that glycosylation is promoting a closer association with neighbouring residues.

We next looked at differences in the pairwise interactions over the course of the simulation. Similar to the Ca distances, the N-linked Asn site generally increased the number of persistent interactions with neighbouring residues on glycosylation (Figure S8). With regards to active site residues, the guanidino group of Arg38 increases its propensity to form H-bonds with Gly69 and Ser73 (Figure 4e). Asn70 likewise, increases its van der Waals interaction network with spatially local residues. Phe41 on the other hand has a lower propensity to form a pi-pi stacking interaction with Phe152 on glycosylation. A pi-pi interaction with Phe152 in the non-glycosylated form could potentially disrupt the conformation of Phe41 relative to haem and thus may impact negatively on catalysis.

Structure-based calculations can excel in detecting networks of rigid clusters through proteins and we have successfully used the FLEXOME implementation of pebble game rigidity analysis previously for this purpose^42,50^. We initially performed rigidity analysis on the available holo-HRP structure^19^. The main change in rigidity occurs as the cut off is reduced from −1.0 to −3.0 kcal/mol over steps of 0.5 kcal/mol (Figure 5a and Figure S10). When the cut off is small, so that even relatively weak hydrogen bonds are included as constraints, a single large rigid cluster extends across almost the entire structure. As the cut off becomes more negative, excluding the weaker hydrogen bonds, peripheral portions of the structure become flexible, while the central part of the protein around the haem group retains rigidity. At the lowest cut offs explored here (−3.0 to −4.0 kcal/mol), the largest rigid cluster includes only about twenty residues around the haem group (Figure 5e), representing the “rigid core” of the protein.

**Figure 5.**
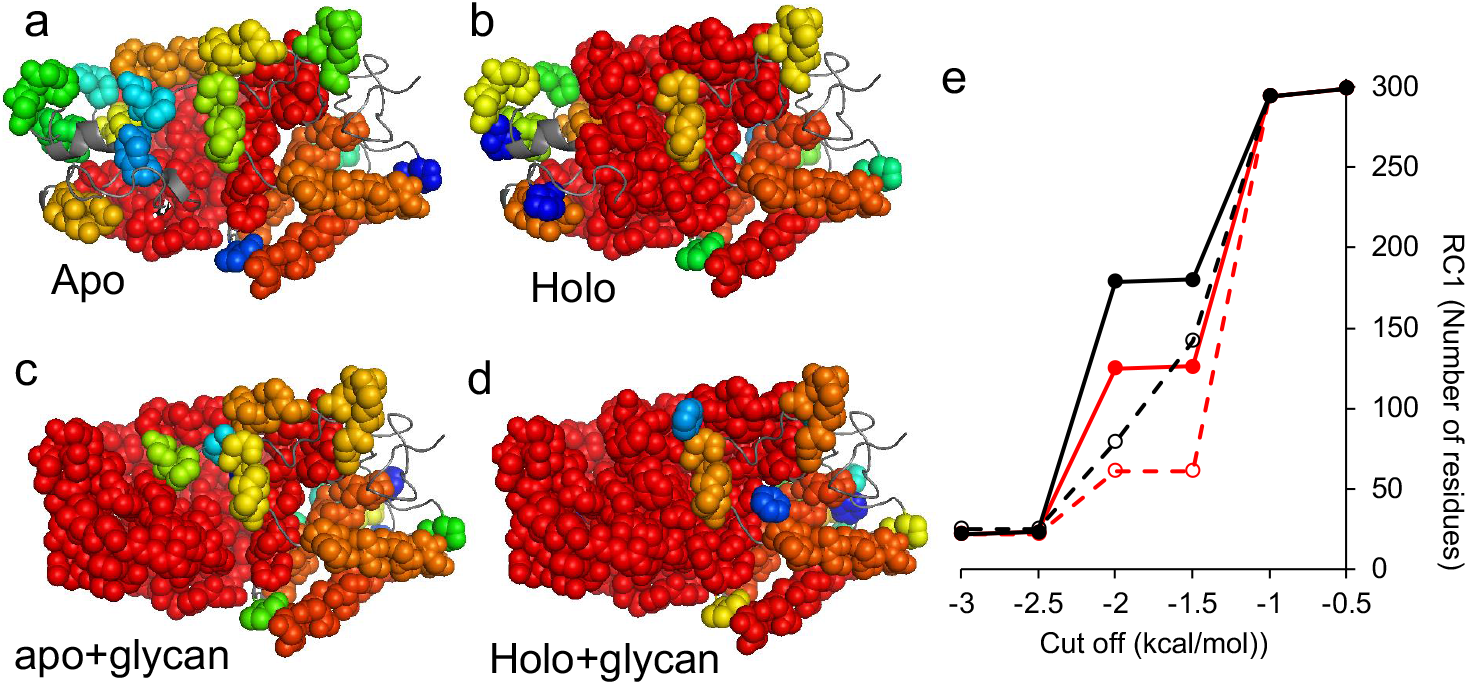
Rigidity analysis of the different HRP forms. (a-d) Structural representation of hydrogen-bond energy cut off of −1.5 kcal/mol for each form. The twenty largest rigid clusters are shown in space-filling representation and rainbow-coloured from red to blue. Flexible regions are shown as grey cartoon. (e) Cluster size at each cut off point. RC1 value is the number of residues which are members of the largest rigid cluster. Red and black represent unglycosylated and glycosylated forms, respectively, with dashed and full lines representing the apo- and holo-forms, respectively.

As there are no structures available for the apo-protein or glycosylated HRP, we used the available X-ray crystal structure for holo-rHRP to produce models. The apo-HRP was generated by simply removing haem from the structure coordinate file prior to analysis. While our experimental evidence shows apo-rHRP is unlikely to have any discernible structure, the cluster analysis does allow us to demonstrate the effect of haem removal on the native structure. Removal of haem has a substantial effect on rigidity of the protein (Figure 5b,e) as haem itself is a rigid body and forms a large number of non-covalent interactions, mostly hydrophobic interactions with surrounding residues. As a result, apo-HRP loses its rigidity much more rapidly as the cut-off becomes more negative. At a −1.5 kcal/mol cut off, only 61 residues comprise the largest rigid cluster for apo-HRP compared to 126 for holo-HRP. This confirms experimental data regarding the importance of haem to maintaining the structural integrity of HRP.

To generate a rigidity mimic of the glycosylated protein, based on previous observations^51–54^ we assumed that attachment of a bulky glycan to the modified asparagine residues will reduce conformational flexibility in the vicinity of that residue, which is largely borne out in the MD simulations (Figure 4a). We therefore introduced additional artificial constraints in the bond network between the Ca of the glycosylated residue and the Ca of the preceding and following residues in the sequence. The additional constraints are added to the bond list before rigidity analysis is carried out. These additional constraints rigidify both the apo and holo forms with respect to non-glycosylated form (Figure 4). As in the case of the apo/holo comparison, the difference is observed in the intermediate cut off regime (−1.5 and −2.0 kcal/mol; Figure 4e).

From Figure 5, we find that rigidification due to glycosylation is visible in peripheral regions of the protein, whereas removal of the haem group has more effect in the central region around the haem binding pocket. That is, our rigidity analysis shows a logical finding that PTMs affect rigidity local to the site of the PTM. However, rigidity is not a strictly local phenomenon but rather percolates across constraint networks; we see in Figure 5 the extent of rigidity in the glycosylated HRP structures, such that peripheral and central regions form a single rigid cluster, whereas in the non-glycosylated HRP structures the periphery is flexible relative to the central region. As a global view we find that the trend in rigidity, based on the size of the largest rigid cluster in the intermediate cut off regime, is Holo-pHRP > Holo-rHRP > Apo-pHRP > Apo-rHRP. This trend exactly mirrors the CD stability data from the extracted *T*_m_ values (Table 1) and the REES data (Figure 2), acknowledging the evident unfolding of Apo-rHRP.

## Discussion

PTM events such as glycosylation and co-factor binding are important in biology by imparting essential structural and functional features on a protein. We show here that these two PTM events also influence protein dynamics leading to enhanced stability, and in the case of glycosylation, improved catalytic performance. Our data shows remarkable correlation between protein stability and active site rigidity. That is, the ranked rigidity of the active site from the Trp117 REES probe is Holo-pHRP > Holo-rHRP > Apo-pHRP >> Apo-rHRP, which mirrors the CD stability data. Our data suggests that glycosylated HRP is a better enzyme both in terms of stability and activity compared to the equivalent non-glycosylated form, with both experimental and simulation data indicating rigidification of the active site playing a major role.

Co-factors such as haem commonly become integral structural components essential to protein structure and stability as well as being necessary for function^2^. This holds true for HRP, with the haem co-factor being buried within the enzyme’s structure making extensive interactions with the protein. Removing haem therefore has an adverse effect on stability and structure as whole, as our data clearly shows (Figures 1–3). Glycosylation on the other hand occurs on the surface of a protein at multiple sites distant from the active site of enzymes, as illustrated here with HRP (Figure 1a); the closest N-linked site is ~20 Å away from the central iron atom of haem. Thus, the effect of glycosylation on HRP, especially the turnover enhancement, is not immediately obvious. We propose that glycosylation of HRP has a dual effect linked by a common mechanism: improved stability and activity through rigidification.

The role of glycosylation in protecting against events such proteolysis and aggregation together with improving solubility is well established and are largely considered the main mechanisms for maintaining a correctly folded and active protein^3^. Improved thermodynamic stabilisation, as observed here for HRP and for other glycoproteins (see Herbert et al^52^ and Sola and Griebenow^55^ for an overview) is also an important contribution. Indeed, our results show that glycosylation can in part make up for the loss of haem by producing a more helical structured and stable apo-HRP (Figures 2–3); this is despite none of the N-linked sites in HRP residing in a helix and previous work suggesting glycosylation rarely induces helical structure^56^. Thus, glycosylation may have a chaperone effect assisting HRP folding; glycans assisting folding has been proposed previously^7^. Increased thermostability can be achieved by reducing the entropic contribution to the free energy of folding through for example, reducing the flexibility of the folded protein. Reduced flexibility does appear to be the case here with glycosylated HRP as demonstrated initially through the REES effect (Figure 3) and backed up with molecular dynamics (Figure 4).

While glycosylation stabilises HRP, another direct benefit of increased rigidity of glycosylated holo-HRP is a 10-fold enhancement of OPD turnover (Figure 2 and Table 1). Enhancement of activity upon glycosylation has been observed for various enzymes^5, 52^ but little is known how glycan addition at sites remote from the active site are influencing enzyme activity. Indeed, it has been observed previously for HRP when plant produced enzyme is compared to the recombinant form^16,21, 28,57^ but with little attention paid and no rationale provided for such an observation. Thus, the main cause of this activity enhancement is largely unknown. One of the main ideas is that general stabilisation of a protein leads to a higher population of correctly folded active enzyme on glycosylation therefore a higher observed activity. However, this is unlikely to be the single major factor for HRP as the unglycosylated protein is structurally stable over the optimal activity temperature (Figure 2a) and has a turnover temperature optimum like that of the glycosylated form (Figure 2b). Moreover, while *k*_cat_ changes *K*_M_ is similar for both holo-HRP forms suggesting changes to the catalytic process or the population of HRP capable of undertaking catalysis at any specific time point. Both our experimental (Figure 3) and simulation (Figure 4–5) data suggest that enhanced activity is likely to be driven by propagated dynamical changes on glycosylation that lead to rigidification of the active site.

Active site dynamics have emerged as having a significant effect on enzyme turnover, and particularly manifesting in a measurable heat capacity of catalysis^10, 58^. Relevant to this study, recent work has shown that chemically induced structural rigidification of an artificial haem peroxidase leads to increased helical content and turnover^33^. Unlike chemically induced rigidification, glycosylation is highly specific in terms of the structural regions directly impacted.

Indeed, the site of N-linked glycosylation is as critical as the glycan addition^52^; glycosylation at engineered non-native sites will not automatically lead to enhanced stability and function^59, 60^. Additionally, significant local rigidification of the N-linked site on glycosylation does not appear to be universal for HRP according to our MD simulations (Figure 4a). Protein engineering has shown that mutational events can change conformational dynamics that in turn shape enzyme activity^47,61–63^. But this is different to glycosylation as protein engineering changes the inherent amino acid sequence itself, commonly through mutations in or close to the active site, not the intrinsic dynamics of the existing protein sequence. Also, given the significant proportion of secreted proteins that are enzymes, glycosylation does appear to be a common mechanism nature uses to mediate enzyme activity.

So how does glycosylation at surface residues distant form the HRP active site impart their effect on activity? Given that the REES probe, Trp117, is close to the active site (Figure 3a), we can assume that the active site becomes more rigid. The experimental data is backed up by the molecular dynamic simulations (Figure 4) and rigidity analysis (Figure 5), with the former showing clearly that residues comprising the active site pocket and substrate tunnel being more rigid in the glycosylated form. Moreover, pairwise analysis shows that Ca are more often closer together in the glycosylated form, with key catalytic residues forming more persistent interactions (Figure 5d-e). Glycosylation may also cause less favourable interactions that may impact on catalysis to become less persistent, such as the pi-stacking interaction between Phe41 and Phe152 (Figure 5e). MD also suggests it may not always be the nearest N-linked glycosylation site in terms of the amino acid sequence that exerts the greatest effect; the rigidification of active site pocket (residues 130-148) is likely to be derived from long range interaction changes associated with N-linked glycosylation of N255, the residue that undergoes the largest reduction in Ca flexibility on glycosylation (Figure 4a). Thus, significant local rigidification of selected N-linked sites and associated propagated network changes may be enough for both enhanced function and thermal stability. Future work that involves systematic mutation of each N-linked site followed by native glycosylation in a plant system could address this but is currently technically quite challenging given *A. rusticana* would need to be the ideal production host to preserve evolutionary-linked glycosylation patterns and isoform production. What it does show is that currently the native enzyme from *A. rusticana* is the best source of HRP in terms of stability and activity towards OPD compared to its sequence equivalent recombinant enzyme. However, given that our work suggests that HRP activity and stability is linked to inherent enzyme rigidity, protein engineering could be used accomplish the same task, with some notable successes providing inspiration.^21, 57,64–66^

To conclude, glycosylation of HRP does not simply appear to be an inert modification protecting the enzyme from aggregation but affects global protein stability and flexibility to the level of altering active site rigidity and thus function. Rigidification of HRP is correlated with a 10-fold increase in turnover but that this rigidification does not affect the temperature optimum for catalysis. Thus, the temperature optimum of catalysis and overall temperature stability of HRP are not necessarily directly correlated, coupled or the same. Glycosylation thus provides HRP with both a functional and stability advantage.

## ASSOCIATED CONTENT

**Supporting Information**. The following files are available free of charge.

Supporting Figures 1-9 and Supporting Table 1 (file type, i.e., PDF)

## ACKNOWLEDGMENT

We would like to thank the Protein Technology Hub facility in the School of Biosciences, Cardiff University for access to protein purification and analysis facilities.

## ABBREVIATIONS

CD, circular dichroism; CSM, centre of spectral mass; MD, molecular dynamics; pHRP, plant horseradish peroxidase; rHRP, recombinant horse radish peroxidase; REES, Red edge excitation shift; RMSF, root mean squared fluctuation.

## Supporting Information

**Figure S1.**
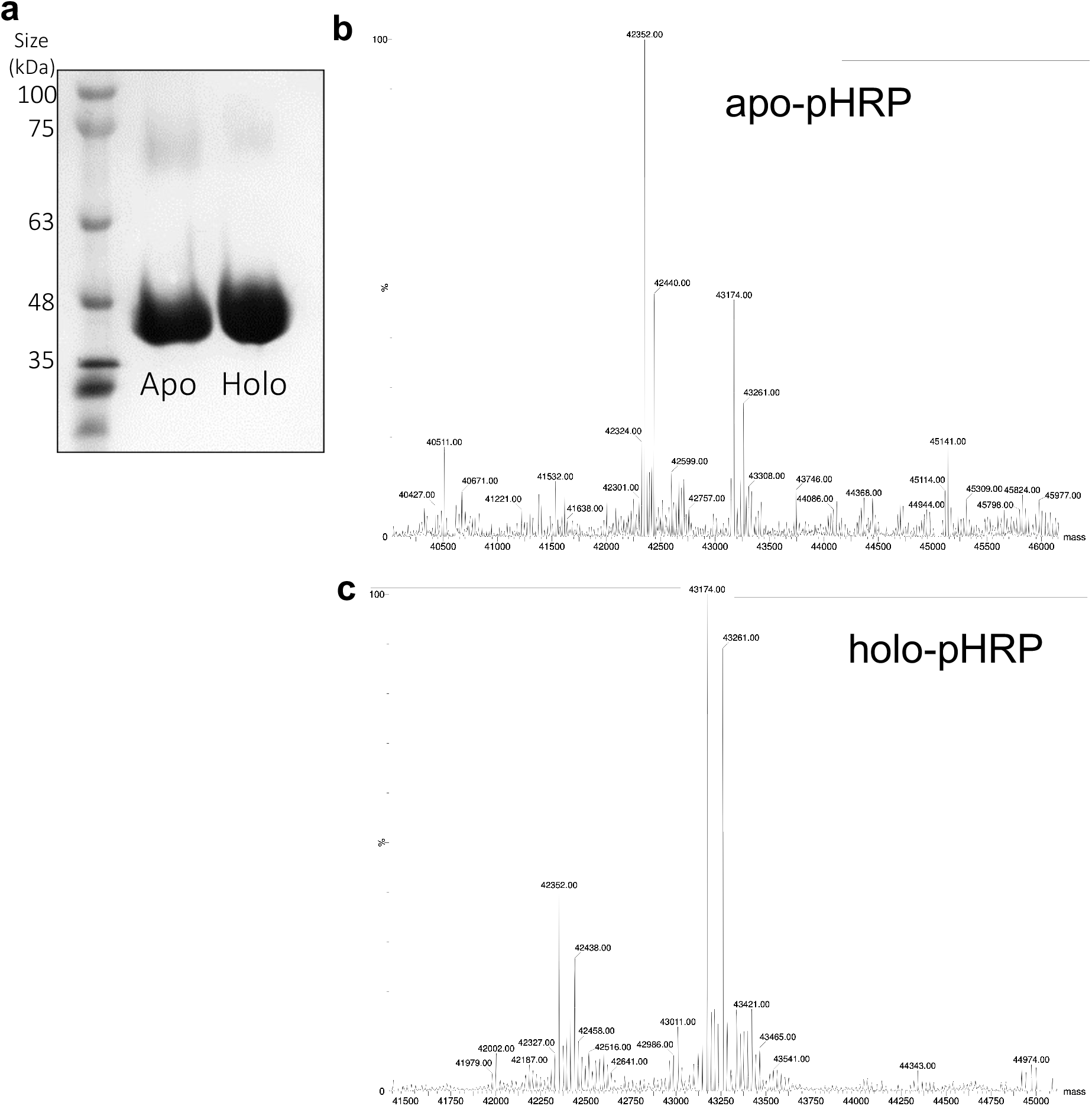
Analysis of the plant HRP. (a) SDS PAGE and (b-c) mass spectrometry of (a, b) apo and (a, c) holo plant HRP. **Supplementary analysis.** The pHRP samples are directly used in immunoassays manufactured by OrthoClinical Diagnostics (OCD). The apo-rHRP is generated by an organic extraction process by the supplier and used as a correction agent in certain OCD diagnostic immunoassays. SDS PAGE confirms the samples’ purity. Mass spectrometry was used to analyse the composition of both the apo (b) and holo (c) versions. Previous studies have shown that the main polypeptide unit is comprised of Gln31-Ser338, which has a theoretical mass of 33918 Da^1–2^. Both samples show two main peaks at ~42kDa (42352 Da and 42440 Da) and ~43 kDa (43174 Da and 43261 Da), with each divided into 2 sub peaks. The masses match previously analysed plant HRP samples^3^. The difference in mass between the two sub peaks is 87-88 Da, which is equivalent to the processing of the C-terminal serine, a common feature of the C1A isoform^3–5^. The two main glycan forms have an estimated masses of 8530 Da and 9351 Da. They represent 7 N-linked carbohydrate units (GlcNAc-2, Man-3, Fuc-1, Xyl-1) with one disaccharide (GlcNAc-1, Fuc-1) and 8 N-linked carbohydrate units (GlcNAc-2, Man-3, Fuc-1, Xyl-1), respectively, as reported previously^3, 5^. Both apo and holo forms of pHRP contain the same components but with the apo-form having an apparent higher proportion of the single disaccharide glycan form. The haem is non-covalently bound so will be lost during the measurement process; thus both measured masses will be equivalent to the apo-protein mass.

**Figure S2.**
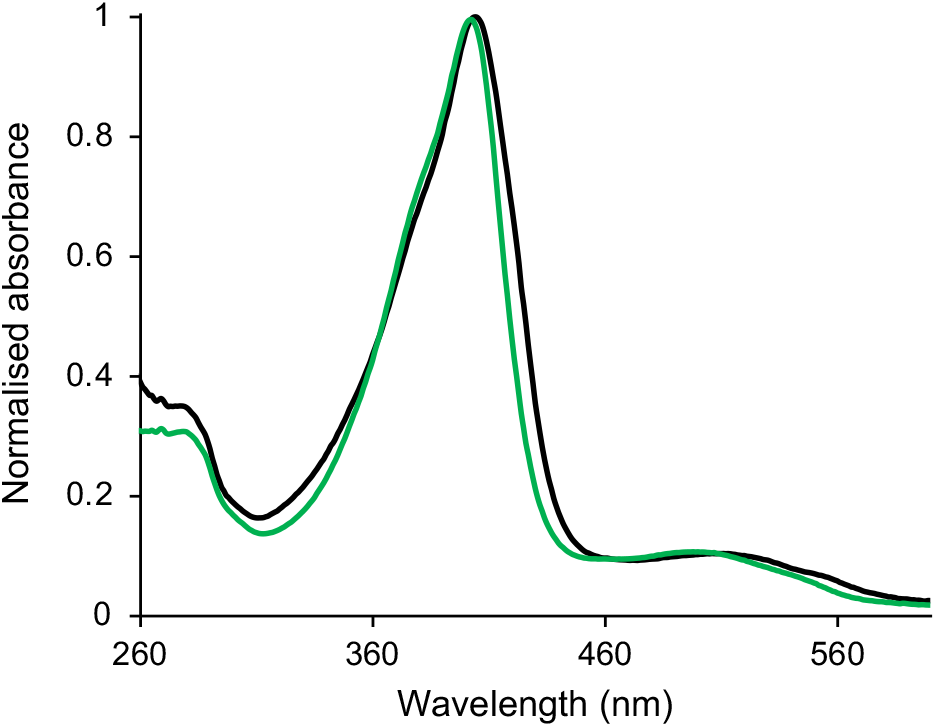
Absorbance spectra of holo-pHRP (green) and holo-rHRP (black).

**Figure S3.**
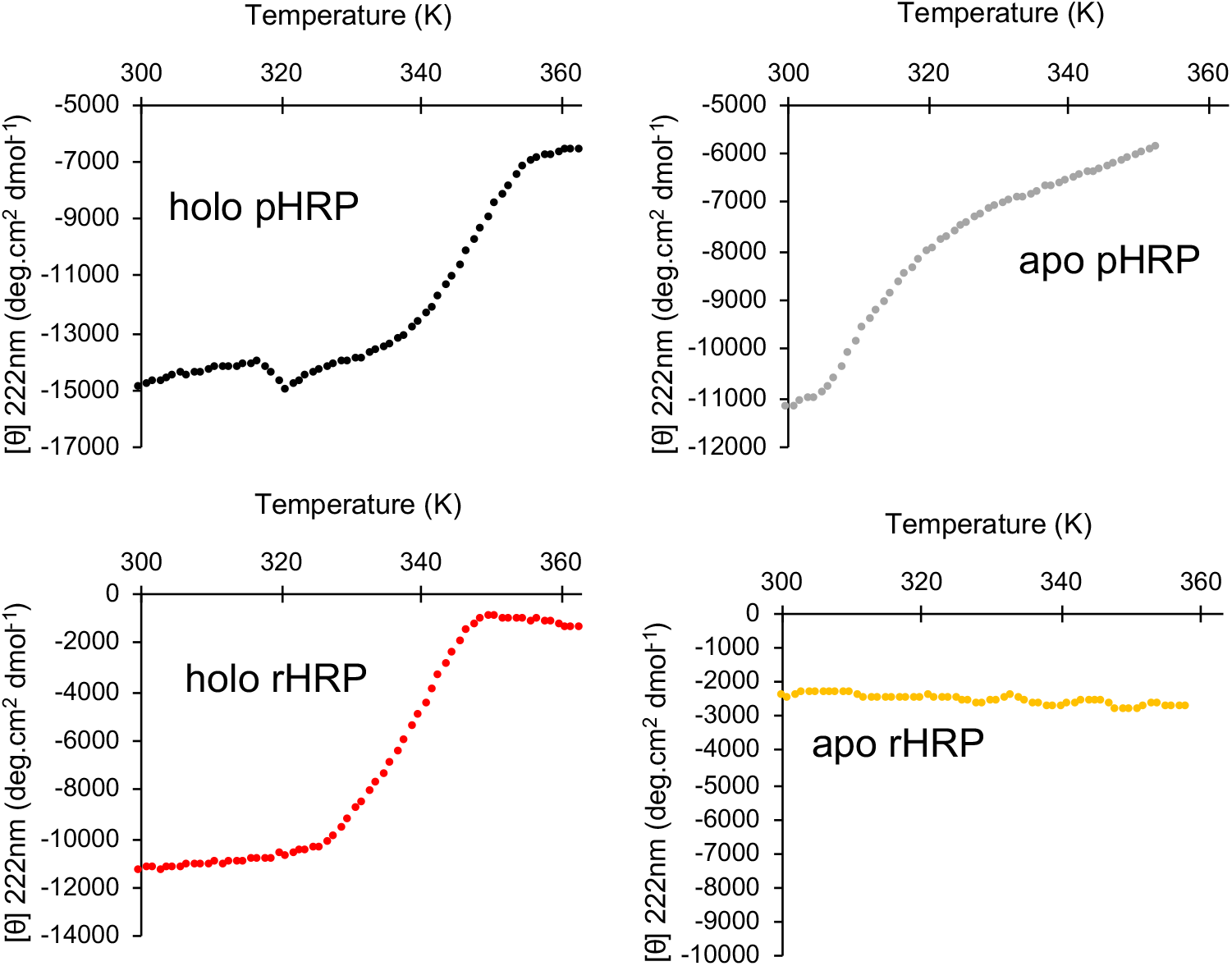
Thermal melt data as measured by CD at 222nm for each form of HRP.

**Figure S4.**
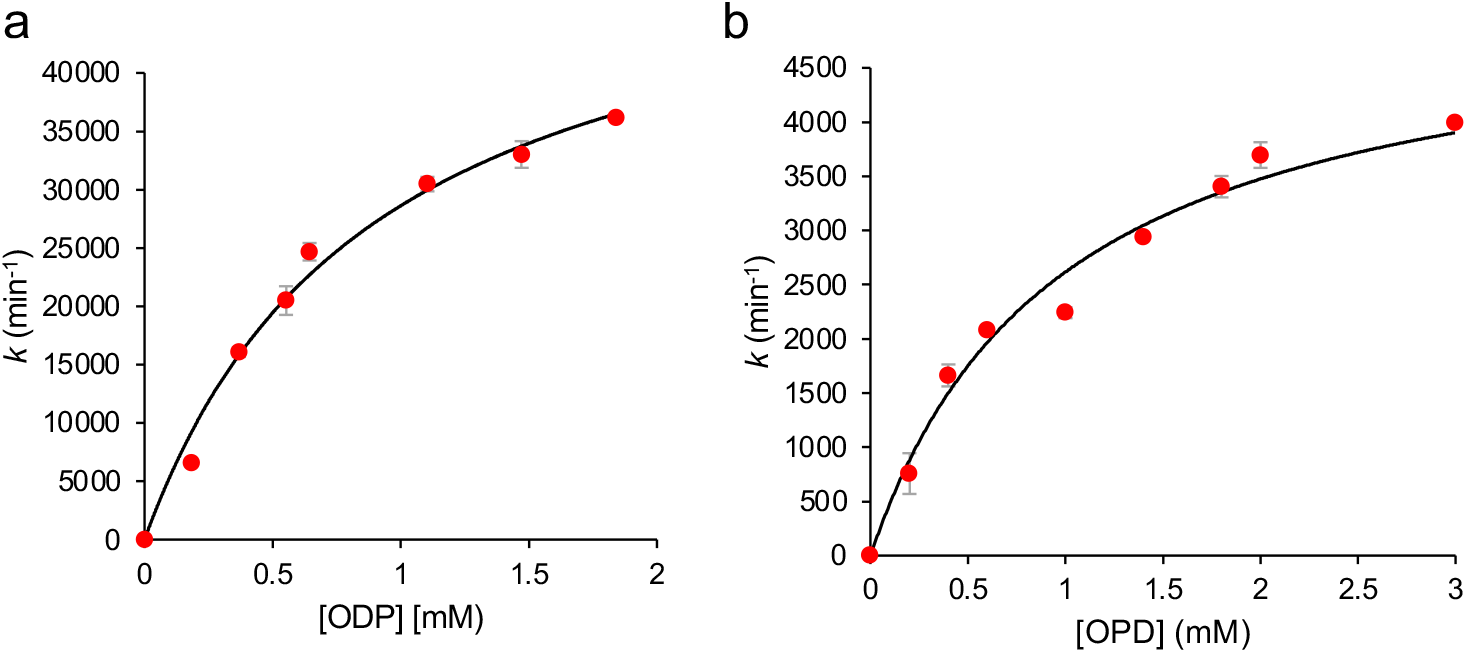
Steady state enzyme kinetics of (a) holo-pHRP and (b) holo-rHRP. The rate was determined by monitoring the production of DAP at 450 nm from OPD. The rates where fit to the Michaelis Menten equation.

**Figure S5.**
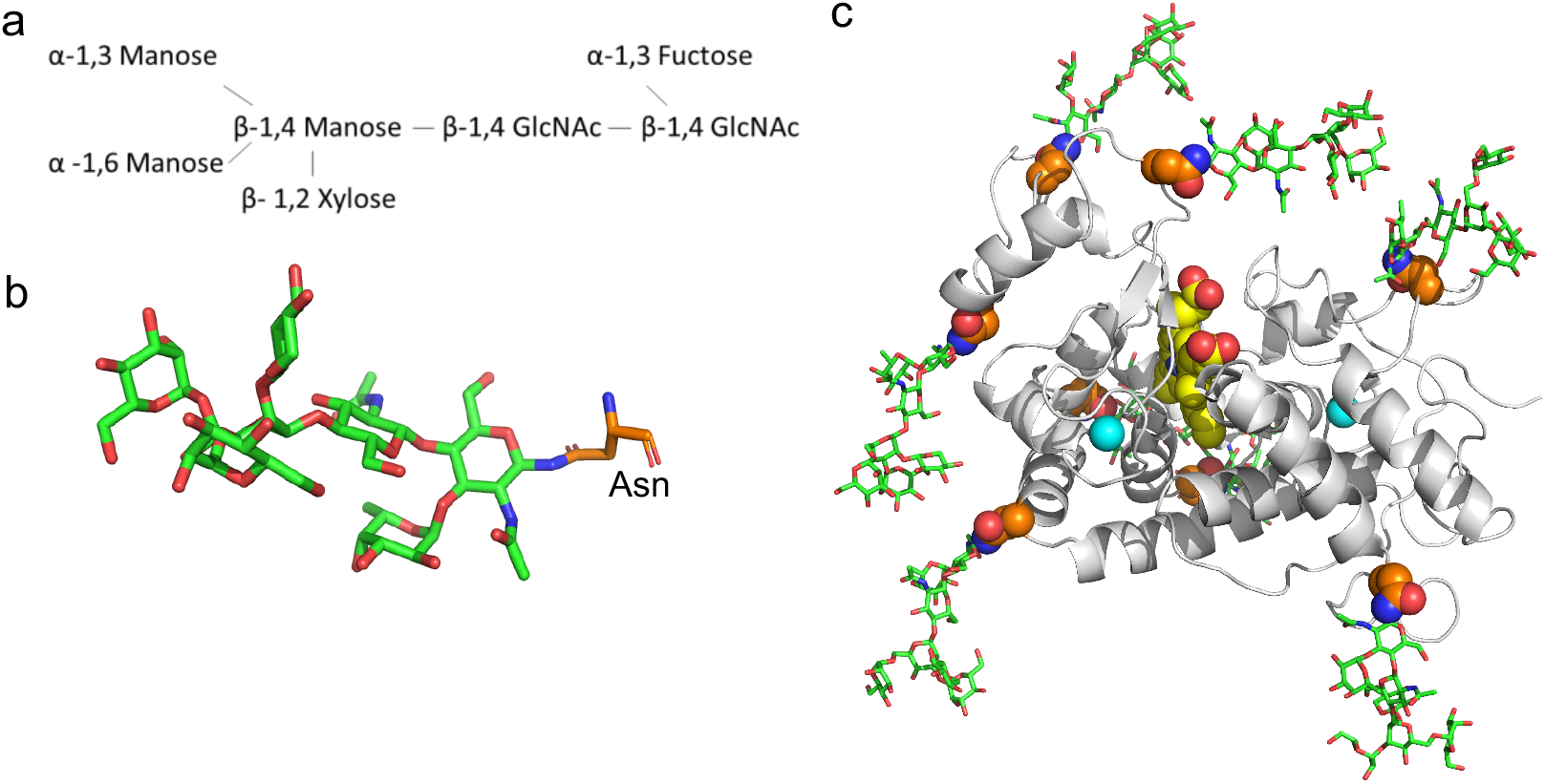
Glycosylated model of HRP. (a) Outline of the glycan composition and connectivity together with (b) a representative N-linked unit bound used in the simulation. (c) Clustered average of the glycosylated HRP model with the glycan units coloured green, the asparagine residues orange, calcium cyan apheres, and haem as yellow spheres.

**Figure S6.**
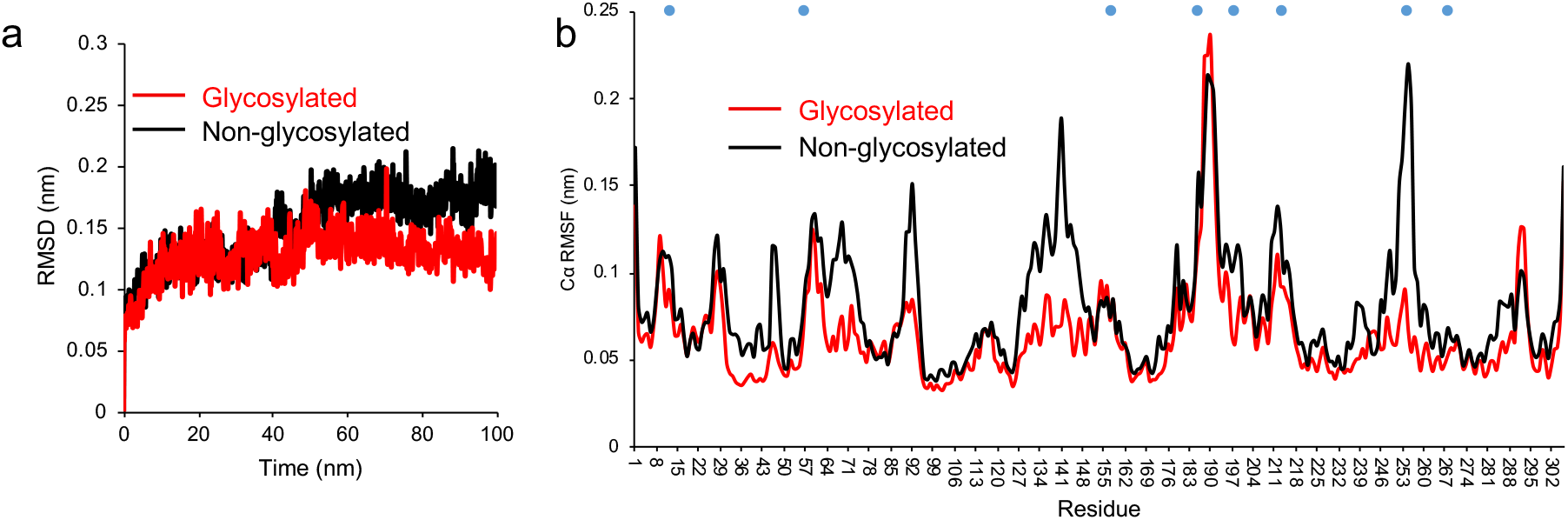
Molecular dynamics of glycosylated (red) and non-glycosylated (black) HRP. (a) Root mean squared deviation (RMSD) and (b) Ca root mean squared fluctuation over a 100 ns simulation. The blue circles represent the N-linked glycosylation sites.

**Figure S7.**
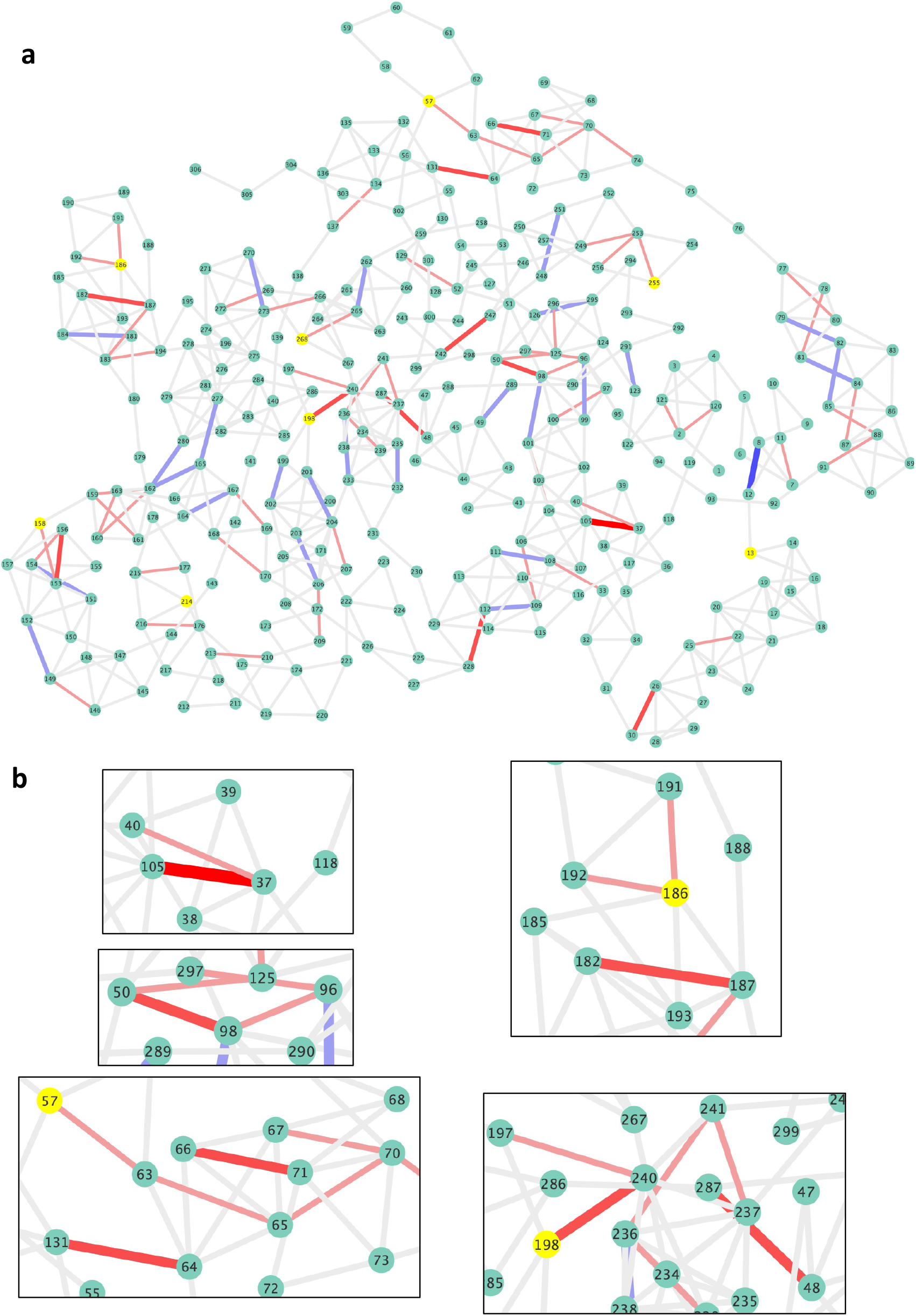
Pairwise network of Ca atoms with a 5.5Å cut-off over the course of the MD simulation. Grey links represent no change in the Ca pairwise distances with red lines indicating closer distances in the glycosylated form and blue lines indicating closer distances in the non-glycosylated form with shade of colour and thickness of the line related to the change in distance (ranging from light colour thinner lines representing ~0.77Å to thicker darker red lines representing up to 1.15Å). (a) the whole pairwise network and (b) selected regions. N-linked glycosylation residues are shown a yellow spheres.

**Figure S8.**
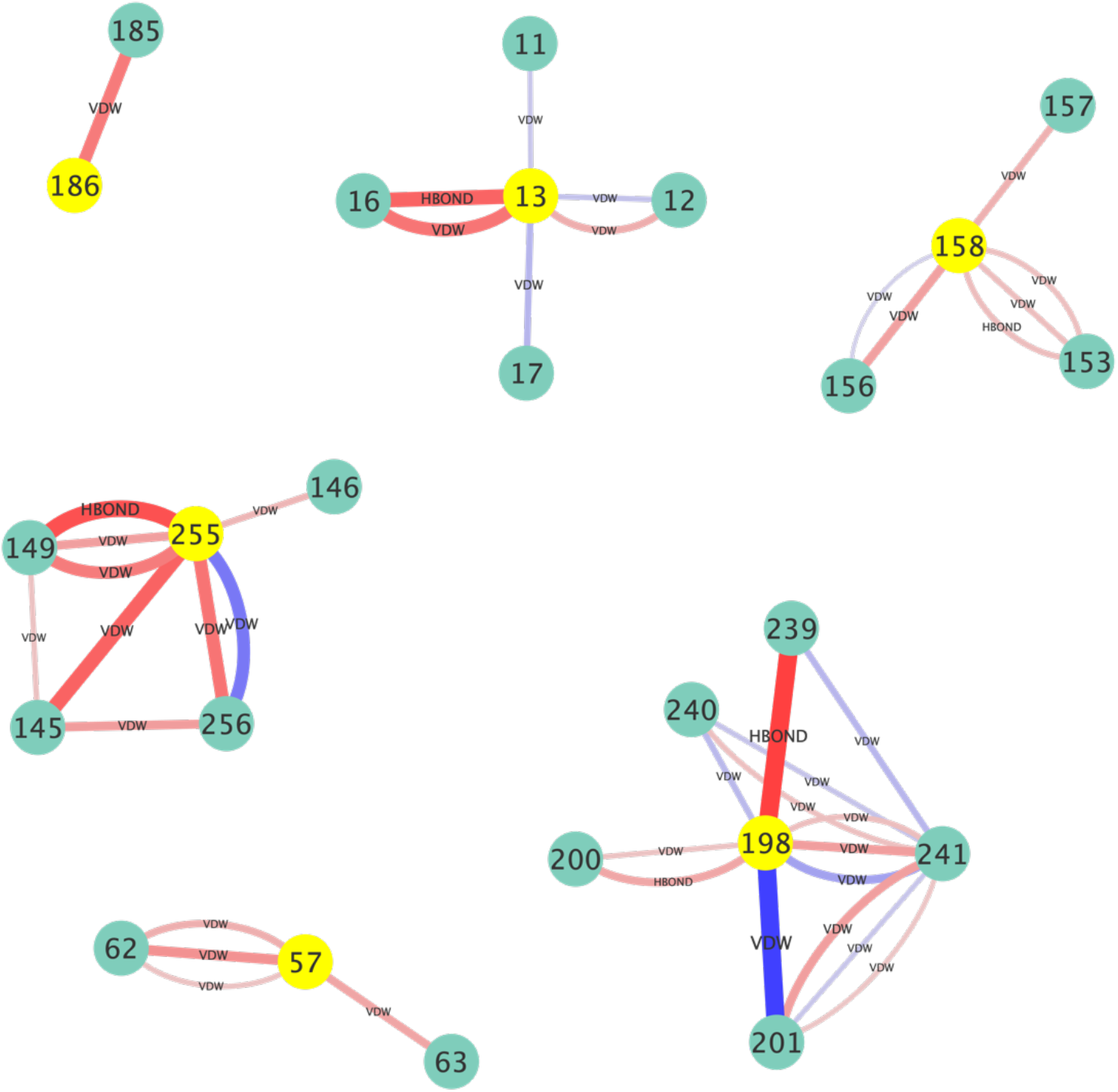
Change in pairwise interactions related to N-linked glycosylated Asn residues (yellow spheres) over the course of the MD simulation. Red lines indicate increased number of interactions in the glycosylated HRP and blue lines increased number of interactions in the non-glycosylated form. The thickness of the lines corresponds to the frequency of an interaction over the course of the MD, with thicker lines representing more persistent interactions. An arbitrary 10% cut off was applied, with interactions differences below this value ignored. The change in interaction type is shown on the diagram with VDW and HBOND equivalent to van der Waals and H-bonds, respectively.

**Figure S9.**
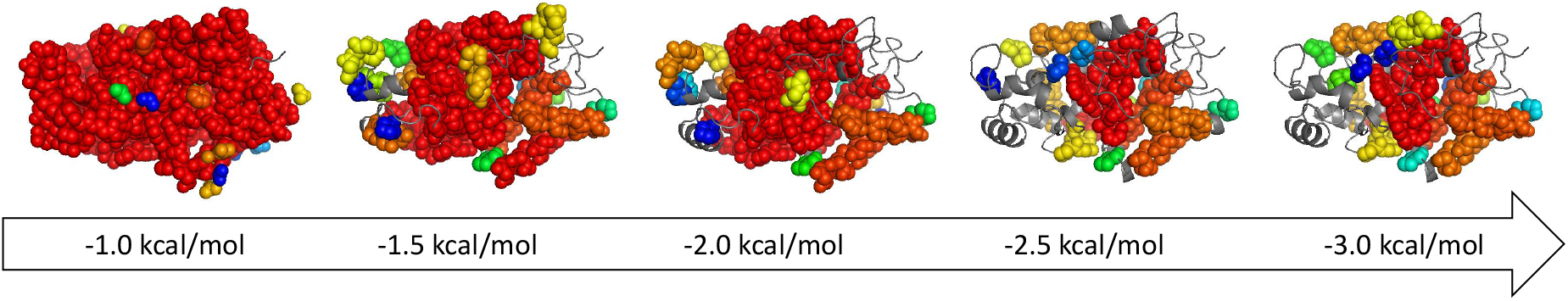
Rigid Cluster Decompositions (RCDs) of holo-HRP at different hydrogen-bond energy cut-offs in the course of a rigidity dilution. The twenty largest rigid clusters are shown in spacefilling representation and rainbow-coloured from red to blue. Flexible regions are shown as gray cartoon. The largest rigid cluster, RC1 (red), spans almost the entire structure (at left) when the cutoff is small. Outlying portions of the structure become flexible as the cutoff becomes more negative during the dilution, until the largest rigid cluster includes only about twenty residues surrounding the haem group (at right). The dilution thus reveals the relative rigidity of different portions of the structure.

**Table S1.** Rigidity analysis of the different HRP forms.^a^RC1 value is the number of residues which are members of the largest rigid cluster. The size of the largest rigid cluster for holo- and apo-HRP with and without glycosylation is shown in the course of a rigidity dilution as the hydrogen-bond energy cutoff is lowered from −0.5 to −4.0 kcal/mol. All structures are almost entirely rigid at small cutoffs (−0.5, −1.0 kcal/mol) and mostly flexible at large cutoffs (−2.5 to −4.0 kcal/mol). Differences in the relative rigidity of the structures, due to the presence/absence of haem and glycosylation, are visible at intermediate cutoff values (−1.5, −2.0 kcal/mol).

| Cut off<br>(kcal/mol) | RC1 value<sup>a</sup> |  |  |  |
| --- | --- | --- | --- | --- |
|  | Holo | Apo | Holo+glycan | Apo+glycan |
| -0.5 | 299 | 299 | 299 | 299 |
| -1.0 | 294 | 294 | 294 | 294 |
| -1.5 | 126 | 61 | 180 | 142 |
| -2.0 | 125 | 61 | 179 | 79 |
| -2.5 | 23 | 22 | 23 | 25 |
| -3.0 | 22 | 22 | 22 | 25 |
| -3.5 | 16 | 16 | 25 | 25 |
| -4.0 | 16 | 16 | 18 | 18 |

---

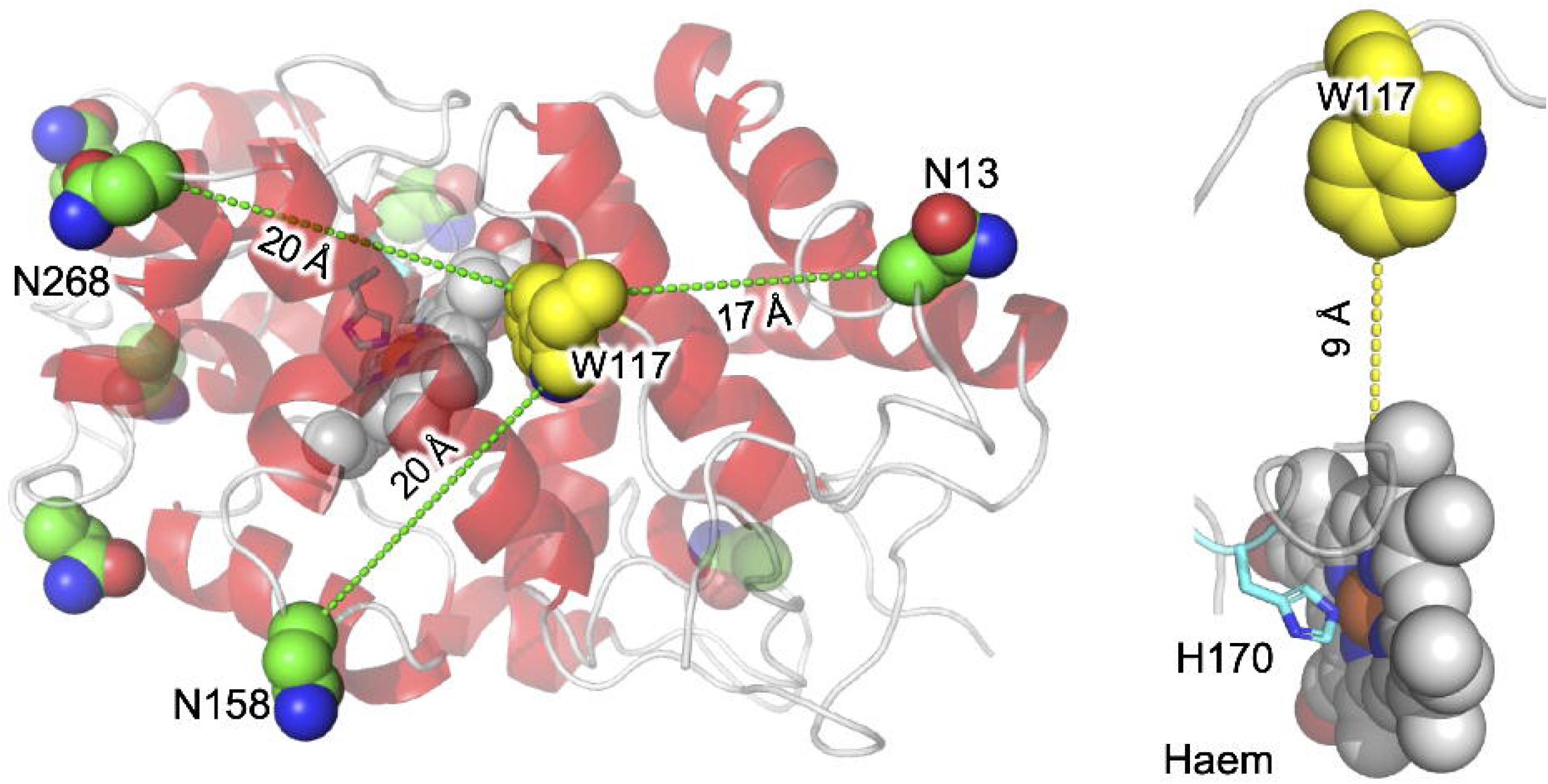

